# Triploidy affects standard and postprandial metabolism in brook charr, *Salvelinus fontinalis*

**DOI:** 10.1101/2020.07.30.229823

**Authors:** Nicole J. Daigle, Charles F.D. Sacobie, Christine E. Verhille, Tillmann J. Benfey

## Abstract

The use of sterile triploids in aquaculture is currently limited because of reduced performance in situations of aerobic stress such as high temperature, hypoxia, and exhaustive exercise. Many studies have therefore attempted to find underlying metabolic differences between triploids and their diploid counterparts to improve triploid rearing protocols. This study investigated the effects of triploidy on postprandial metabolism (and therefore also pre-feeding standard metabolic rate; SMR) by measuring oxygen uptake and total ammonia nitrogen (TAN) excretion at 14-15°C of previously fasted (for eight days) diploid and triploid brook charr, *Salvelinus fontinalis*, from 48h before to 48h after being fed a single ration of 0.4% body mass. Triploids had significantly lower SMRs and higher postprandial metabolic rates (i.e., specific dynamic action) and net TAN excretion than diploids. While this greater cost of processing a meal may not represent a major diversion of metabolic reserves for triploids, it could affect their growth and survival when simultaneously faced with oxygen-limiting conditions.

## 1. Introduction

Triploids contain three complete sets of homologous chromosomes in their genomes, resulting in reproductive sterility in most species of fish (Benfey 1999; Piferrer *et al.* 2009). This aspect of their biology makes the use of triploids a practical approach for preventing genetic introgression of maladaptive alleles from escaped farmed fish into wild populations (Benfey 2016), which is an issue of great concern for Atlantic salmon (*Salmo salar*) conservation (Glover *et al.* 2020). Furthermore, flesh quality is not diminished in triploid females by the repercussions of sexual maturation (Manor *et al.* 2014). Since very little energy is allocated to gonadal development in triploid females, the metabolized energy can instead be put towards somatic growth, although this is not always realized (Piferrer *et al.* 2009).

Despite these advantages, triploidy has yet to become widely adopted in salmon aquaculture because in stressful environments (e.g., high temperature, hypoxia, and exhaustive exercise), triploids perform poorly compared to their diploid counterparts (Hyndman *et al.* 2003a; Hansen *et al.* 2015; Sambraus *et al.* 2017a; Sambraus *et al.* 2018). Therefore, many studies have attempted to find physiological differences between triploids and diploids, as pinpointing such differences could allow for aquaculture practices to compensate for them. For instance, the recognition that nutritional requirements differ between ploidies of Atlantic salmon (Taylor *et al.* 2015; Smedley *et al.* 2016) led to the reformulation of standard diploid diets to reduce the prevalence of ocular cataracts and skeletal deformities in triploids (Sambraus *et al.* 2017b; Peruzzi *et al.* 2018).

A possible explanation for why triploidy influences performance is that fundamental changes in their cell size and number (Benfey 1999) affect metabolic processes under suboptimal conditions to a greater extent than in diploids. For instance, while numerous studies have not found a significant effect of triploidy on standard or routine metabolic rates (SMR and RMR, respectively) at or near optimum temperatures for diploids (Sezaki *et al.* 1991; Parsons 1993; Hyndman *et al.* 2003b; Bernier *et al.* 2004; Lijalad and Powell 2009; Bowden *et al.* 2018), experiments conducted at higher temperatures suggest that triploids have a lower thermal optimum for metabolism than diploids (Atkins and Benfey 2008; Sambraus *et al.* 2017a; Sambraus *et al.* 2018). Routine metabolic rate is the metabolic rate of a fish in a post-absorptive state that is experiencing routine (but minimal) periods of activity; RMR is distinguished from SMR in that measurements taken during minor activity are included, whereas SMR involves protocols to exclude periods of significant activity (Chabot et al. 2016a). Of direct relevance to this study, juvenile triploid brook charr (*Salvelinus fontinalis*) had significantly lower RMRs than diploids at 13-14°C (Stillwell and Benfey 1996) but higher average SMRs than diploids at 16-17°C (although this was not statistically significant at a *p*-value of 0.05; O’Donnell *et al*. 2017). The optimum temperature for diploid brook charr is 15°C, based on a meta-analysis of laboratory and field studies (Smith and Ridgway 2019).

A less studied aspect of triploid metabolism is specific dynamic action (SDA), which is defined as the amount of energy required to ingest, digest, absorb, and store nutrients/energy (Chabot *et al.* 2016b). Many studies have shown that ration size increases the peak metabolic rate and duration of SDA in ectothermic vertebrates (Jobling and Spencer Davies 1980; Andrade *et al.* 1997; Secor and Faulkner 2002; Fu *et al.* 2005; Jordan and Steffensen 2007). Furthermore, SDA’s peak metabolic rate increases and its duration decreases, with an increase in water temperature in fish (Luo and Xie 2008; Frisk *et al.* 2013), whereas hypoxia has the opposite effect (Jordan and Steffensen 2007; Eliason and Farrell 2014). The only two studies to have investigated SDA in triploids to date found no ploidy-related differences (Oliva-Teles and Kaushik 1987a; O’Donnell *et al.* 2017).

Another important aspect of metabolism is the energetic cost of internal ammonia regulation and its excretion. Oliva-Teles and Kaushik (1987a) found no effect of ploidy on daily ammonia excretion patterns or overall ammonia excretion rates when comparing juvenile diploid, triploid, and tetraploid rainbow trout (*Oncorhynchus mykiss*) and Hyndman *et al.* (2003b) found no significant differences overall in ammonia excretion rates following exhaustive exercise between diploid and triploid brook charr. However, triploids in the latter study excreted large amounts of ammonia within the first hour of exercise, which likely contributed to their significantly faster recovery of plasma osmolality, white muscle lactate, white muscle ATP, and post-exercise oxygen consumption rates. Segato *et al.* (2006) found that triploid shi drum (*Umbrina cirrosa*) had significantly lower retention (storage) of crude protein compared to diploids, implying differences in protein metabolism, and hence ammonia excretion, but the effects of ploidy on ammonia excretion throughout SDA have not been reported.

The objective of this study was to investigate the effects of triploidy on SMR and SDA (by measuring oxygen uptake), and on kidney/gill ammonia excretion (by measuring net total ammonia nitrogen (TAN) excretion) in fish. Brook charr were selected as the study species because they are better suited than Atlantic salmon for rearing in small-scale freshwater research facilities. Both species have similar metabolic physiology and thermal ranges and can be fed the same high-protein diets.

## 2. Materials and methods

### Rearing information

The brook charr used in this experiment were produced through an in-house breeding program (University of New Brunswick, Fredericton) in December 2017. Following fertilization, the eggs were divided into two groups with half retained as diploid controls and the other half exposed to a pressure treatment of 5-min duration at 65.5 MPa (TRC-APVM, TRC Hydraulics Inc., Dieppe, New Brunswick) beginning at 220°C-min post-fertilization. The fish were then reared using standard salmonid aquaculture methods. Fish were size-matched three times prior to experimentation to limit size-variation between ploidies and were given unique identification with passive integrated transponders (PIT-tags) via a 3mm incision made into the lower abdominal cavity (body mass [BM] of 24 ± 0.4g [mean ± SE] at time of tagging). Blood samples were taken during tagging to make blood smears for ploidy confirmation (Benfey *et al.* 1984) based on the length of the major axis for 50 erythrocytes per fish. Using this method, all presumed diploid and triploid fish were confirmed to be of the correct ploidy.

The fish were given approximately two weeks (December 26^th^, 2018 - January 11^th^, 2019) to acclimate to the constant (24h) light and test temperature (14.4 ± 0.1ΰC; mean ± SE; range 13.7-15.0ΰC), during which they were fed 0.6% of tank biomass once daily using 5mm pellets of a salmonid-specific (high protein) diet formulation (Corey Aquafeeds, Fredericton, New Brunswick). Constant light was used to avoid the physiological stress response observed in fish exposed to abrupt on-off light transitions (Ryu *et al.* 2019). This is also consistent with aquaculture practices, as 24h lighting increases growth rates and delays sexual maturation (Harmon *et al.* 2003). The fish were divided among four 68.4L circular tanks (48.8cm diameter, two per ploidy), with an average flow rate of approximately 2.0L/min and a stocking density of approximately 25kg/m^3^. Fish had an average BM of 193 ± 5g (mean ± SE) at the beginning of the trials, which took place from January 11^th^ to March 27^th^, 2019. All experimental protocols were approved by the UNB Animal Care Committee (AUP 18036), adhering to guidelines established by the Canadian Council on Animal Care.

### Experimental trials

Eight days before experimental feeding, two fish per ploidy (four total) were placed into individual intermittent flow-through respirometers (approximately 7.5L each). The sides of the respirometers were covered to minimize visual disturbances to the fish. Individuals were selected from their home tanks using a convenience sampling method based on visual assessment of size (BM, fork length, and condition factor) to ensure that they were appropriately sized relative to the respirometer (approximately 30g BM/L), and then anaesthetized using tert-amyl alcohol (1% solution in fish-culture water) for fork length, volume, and BM measurements. A random number generator was then used to determine both the individual respirometer and the feeding group for each fish.

To get a reliable picture of SDA, fish were fasted in their respective respirometers for eight days then tube-fed a single ration (measured as % BM) (Fu *et al*. 2005; Jordan and Steffensen 2007). To minimize aerobic stress unrelated to feeding, the BM measurements from eight days pre-feeding were used for calculating ration size, SMR, SDA parameters, and net TAN excretion. The average loss in BM over the eight-day acclimation period was subsequently determined to be minimal (2.9%), based on the re-measurement of fish at the end of each trial.

Oxygen measurements were taken continuously from 48h pre-feeding to 48h post-feeding, with 0h defined as the time of feeding, using oxygen sensors (Fibox 4 trace and OXY-1 SMA, PreSens, Regensburg, Germany) that were calibrated before each trial. Background respiration was measured in each of the respirometers (preliminary tests; data not shown) and was considered negligible. The respirometers were on automated timers set to close for 20 min (sealed phase) and then flush for 10 min (open phase) on a repeating cycle. A minimum dissolved oxygen cut-off of 70% air saturation was used in the study to avoid potentially confounding effects of hypoxic stress on the fish (Fu *et al.* 2005). Paired water samples, separated by a 15-min time interval, were taken at 24 and 12h pre-feeding and 2, 4, 6, 7, 8, 9, 10, 11, 12, 24, 30, 36, and 48h post-feeding and frozen at −20°C for later TAN analysis. Initial samples were taken once the respirometer began the seal interval at the pre-determined times listed above and final samples after a 15-min period to ensure adequate time for sampling all respirometers before the flush interval resumed at the 20-min mark.

As pilot studies showed that the fish would not eat voluntarily in the respirometers, they were tube-fed at 0h. This was done by removing each fish from its respirometer, anaesthetizing it (1% tert-amyl alcohol) to the point where it had minimal response to a tail pinch, removing it from the water, placing a food pellet (5mm) at the back of the mouth, and using a ¼-inch diameter piece of wetted flexible tubing to push the pellet into the stomach. This process was repeated until the entire ration was delivered and then the fish was returned to its respirometer for recovery. There were two feeding groups per ploidy in this experiment: a sham-fed control and a 0.4% BM ration (maximum of 5 pellets delivered). The sham-fed fish were handled identically to the tube-fed fish, with the exclusion of pellet insertion. A cut-off time of 1 min out of the water was used for the feeding process to limit hypoxic stress. All fish were closely monitored to recover any regurgitated pellets so that they could be dried and weighed, then have the dry weight subtracted from the total ration delivered. An a-priori decision was made to exclude any fish from the study that regurgitated more than 1 pellet (approximately 0.1% BM).

Fish were video-recorded throughout the trials to monitor their behaviour and to exclude measurements during periods of significant movement (activity above sustained swimming, i.e., erratic behaviour).

At 48h post-feeding, the fish were removed from their respirometers and euthanized (rapid overdose in 10% tert-amyl alcohol) to re-measure their BM as well as the mass of their viscera (for calculation of viscerosomatic index; VSI), liver (for calculation of hepatosomatic index; HSI), and gonads (for calculation of gonadosomatic index; GSI). Visual examination of the gonads at this time was also used to determine the sex of each fish. The cycle of 48h pre-feeding to 48h post-feeding (= 1 trial) was repeated eight times using different individuals for each trial, for a total of 32 fish. However, metabolic rate data were only collected for three of the four fish in each trial (= total of 24 fish) because one of the PreSens oxygen meters had an unacceptably low signal-to-noise ratio. A random number generator determined which respirometer (and thus feeding group) did not have the oxygen levels monitored for each trial.

### Analysis of oxygen uptake

After each trial, mass-specific metabolic rate (MO_2_; mg/kg/h) was determined for each seal interval by calculating the slope of the decrease in dissolved oxygen concentration over the respective time interval, using the formula:

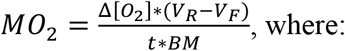

Δ[O_2_] is the change in dissolved oxygen concentration throughout the seal interval (mg/L)
V_R_ is the volume of the respirometer (L)
V_F_ is the volume of the fish (L) measured at the beginning of the trial
t is the duration of the seal interval (h), and
BM is the body mass of the fish (kg) measured at the beginning of the trial

For SMR and SDA calculations, a maximum of 24 and 92 slopes were generated, respectively. Measurements taken during periods of significant movement were excluded from the dataset; therefore, the number of slopes varied based on the activity level of each fish. SMR was determined through quantile analysis of the 12h interval before feeding as this was when the fish were most quiescent (determined by preliminary analysis of the video-recordings). The 0.2 quantiles were used to estimate SMR and SDA, but all SMR quantiles are reported for reference. A general additive model (GAM) was used to determine the predicted model (SDA plot) based on the change in MO_2_ over time post-feeding (visualized in Figure 1).

**Figure 1.**
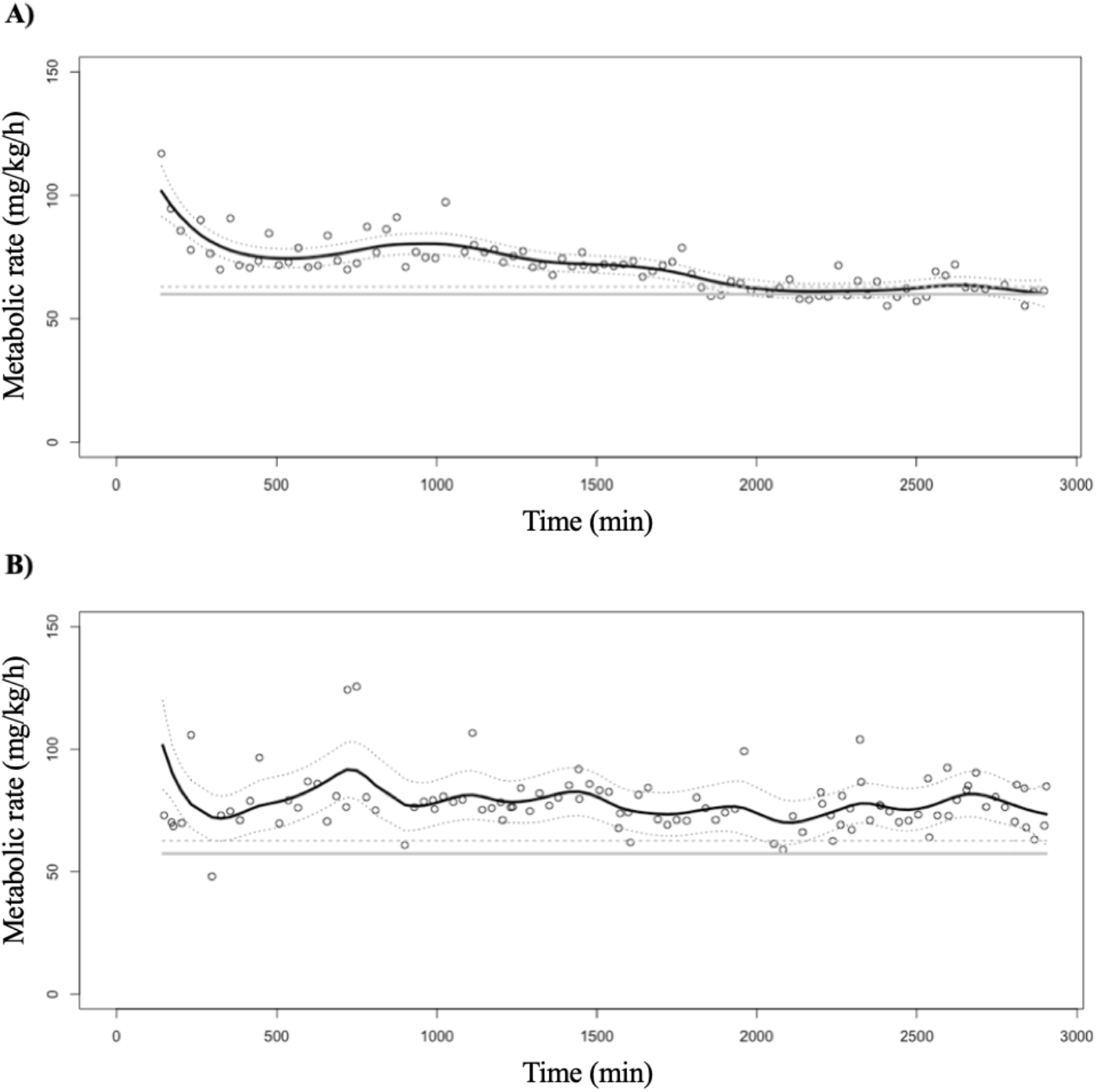
Example of the general additive model for generating the plot for a single **A)** diploid and **B)** triploid brook charr (*Salvelinus fontinalis*) following a designated ration of 0.4% body mass in a single feeding. The predicted model is represented by the black solid line with standard errors shown by the grey dotted lines. Standard metabolic rate (SMR) is represented by the grey solid line with 5% tolerance above SMR shown by the grey dashed line. Each circle on the graph represents a single metabolic rate measurement (i.e., one seal interval). The general additive model used a k-level of 40, “tp” bases (thin plate regression splines), “gamma” family distribution, and SMR was estimated using quantile analysis (0.2 quantiles) from the 12h pre-feeding interval.

Using the fishMO2 (calcSDA function; Chabot 2020) and mgcv (gam function; Wood 2011) packages in RStudio, various SDA parameters were determined for all fish. The actual SDA (referred to as net area; mg/kg) was calculated as the area under the SDA curve (predicted model) bound by SMR for each fish. The duration of SDA (referred to as SDAendtime; min) was defined as the time post-feeding at which the elevated MO_2_ first returned to within 5% of the pre-feeding SMR (standard tolerance value; Chabot *et al.* 2016b). If a fish did not return to this level by 48h post-feeding, the tolerance value was increased to 9%. Tolerance was not increased to 9% for all fish to avoid unnecessarily truncating SDA and thereby decreasing SDAendtime and net area. Peak MO_2_ (referred to as MRpeak; mg/kg/h) was defined as the highest MO_2_ recorded by the predicted model from the interval of 0 to 48h post-feeding. Lastly, the relative (factorial) increase in MO_2_ (referred to as MRfactscope; no units) was calculated as (MRpeak/SMR).

### Analysis of TAN excretion

For TAN quantification, water samples were processed in duplicate using the colorimetric assay outlined by Verdouw *et al.* (1978). Net TAN excretion due to SDA (mmol/kg) was calculated for each fish by first subtracting the average pre-feeding TAN excretion from its TAN excretion at each post-feeding time interval and then adding the values for all time intervals together to create a net TAN excretion value for each fish.

### Statistical analyses

To identify relevant continuous random variables to be included in statistical analyses, preliminary linear regressions among all possible combinations of paired continuous random variables (BM, growth rate, fork length, condition factor, VSI, HSI, and GSI) were completed to identify potential covariates. In the event of two correlated random variables (*p*<0.05), the one that explained the greater amount of variation in the data was selected as representative and the other was removed from the model. Linear regressions were then completed between the remaining random variables and the dependent variables of interest. Continuous random variables that showed significant correlations (*p*<0.05) were included as covariates in the subsequent statistical analyses, as were the categorical random variables respirometer, home tank, trial, and sex.

Effects of feeding group, ploidy, and the interaction between feeding group and ploidy on net TAN excretion, SMR, and SDA parameters of interest (net area, SDAendtime, MRpeak, and SDAfactscope) were investigated using either multiple-factor ANCOVAs (when there were significant covariates) or ANOVAs followed by post-hoc testing to investigate trends, when appropriate. Once the results were generated from each analysis, highly insignificant variables (*p*-value > 0.2) were removed from the statistical model and it was rerun; this process was repeated until the trend was clear (Fisher and Van Belle 1993). If the covariate was significant in the ANCOVA, a post-hoc least squares regression (one-tailed type III z-test) was used to remove the effects of the covariate on the variable of interest.

## 3. Results

Only one of the tube-fed fish regurgitated part of its meal and the amount (one pellet; <0.1% BM) did not meet the criterion for exclusion. However, for the SMR and SDA analyses, 4 of the 24 fish (for which oxygen uptake data were available), were excluded because of poor sensor alignment. This resulted in sample sizes of 4, 4, 6, and 6 for the 2N-Sham, 3N-Sham, 2N-0.4% BM, and 3N-0.4% BM feeding groups, respectively. In five cases, post-feeding MO_2_ did not return to within 5% of SMR, but increasing the tolerance to 9% captured the SDAendtime for three of them. For the remaining two (one diploid and one triploid), the time of the final MO_2_ measurement was used for SDAendtime. For the TAN analyses, no data were excluded, yielding n=7, 7, 9, and 9 for 2N-Sham, 3N-Sham, 2N-0.4% BM, and 3N-0.4% BM, respectively.

Body mass, fork length, and condition factor were not significantly affected by feeding group, ploidy, or their interaction, but VSI, HSI, and GSI were all significantly lower in triploids than in diploids (Table 1). Pre-feeding SMR did not differ between assigned feeding groups (sham versus fed), but all SDA parameters and net TAN excretion values were significantly higher in the 0.4% BM feeding groups than in the sham-fed groups (Table 2; visualized in Figure 2). The variable trial was significant in the analyses of net area (*p*-value 0.014) and MRpeak (*p*-value 0.0021), but the post-hoc testing for these variables did not reveal any significant pairwise trends. A significant covariate effect was found for BM on net TAN excretion and the *p*-value was therefore adjusted through a post-hoc least squares regression (Table 2).

**Table 1.** Results of multiple-factor ANOVAs on the effect of feeding group (FG), ploidy (P), and their interaction (FG:P) on body size parameters (body mass (BM), fork length, and condition factor; measured eight days pre-feeding), and viscerosomatic index (VSI), hepatosomatic index (HSI), and gonadosomatic index (GSI) (measured 2 days post-feeding) in brook charr, *Salvelinus fontinalis* (mean ± SE). For all body parameters, n=7, 7, 9, and 9 for 2N-Sham, 3N-Sham, 2N-0.4% BM, and 3N-0.4% BM, respectively. Statistically significant *p*-values have been bolded (*p*<0.05).

| Variable | 2N-Sham | 3N-Sham | 2N-0.4% BM | 3N-0.4% BM | FG<br>F-ratio,<br>*p*-value | P<br>F-ratio,<br>*p*-value | FG:P<br>F-ratio,<br>*p*-value |
| --- | --- | --- | --- | --- | --- | --- | --- |
| BM (g) | 187.64 ± 7.14 | 186.97 ± 7.31 | 195.4 ± 8.76 | 200.84 ± 13.24 | 0.30, 0.59 | 0.17, 0.69 | 0.091, 0.77 |
| Fork length (cm) | 24.06 ± 0.43 | 24.69 ± 0.43 | 24.87 ± 0.45 | 25.27 ± 0.64 | 1.21, 0.28 | 0.34, 0.57 | 0.048, 0.83 |
| Condition factor | 1.35 ± 0.04 | 1.24 ± 0.04 | 1.27 ± 0.05 | 1.24 ± 0.03 | 2.12, 0.16 | 0.49, 0.49 | 0.87, 0.36 |
| VSI (%) | 11.85 ± 0.46 | 9.17 ± 0.18 | 11.55 ± 0.38 | 9.62 ± 0.24 | 0.41, 0.53 | 18.83, **<0.001** | 1.26, 0.27 |
| HSI (%) | 1.32 ± 0.09 | 1.13 ± 0.07 | 1.45 ± 0.14 | 1.10 ± 0.05 | 0.94, 0.34 | 7.33, **0.011** | 0.74, 0.40 |
| GSI (%) | 3.51 ± 0.73 | 1.49 ± 0.51 | 3.01 ± 0.71 | 1.03 ± 0.40 | 0.35, 0.56 | 6.13, **0.020** | 0.0012, 0.97 |

**Table 2.** Results of multiple-factor ANOVAs or ANCOVAs on the effect of feeding group (FG), ploidy (P), and their interaction (FG:P) on standard metabolic rate (SMR), specific dynamic action (SDA) parameters, and net total ammonia nitrogen (TAN) excretion in brook charr, *Salvelinus fontinalis* (mean ± SE). For SMR and SDA parameters, n=4, 4, 6, and 6 for 2N-Sham, 3N-Sham, 2N-0.4% body mass (BM) and 3N-0.4% BM, respectively, and for net TAN excretion, n=7, 7, 9, and 9, respectively. Statistically significant *p*-values have been bolded (*p*<0.05).

| Variable | Analysis | 2N–<br>Sham | 3N–<br>Sham | 2N–0.4%<br>BM | 3N–0.4%<br>BM | FG<br>F-ratio,<br>*p*-value | P<br>F-ratio,<br>*p*-value | FG:P<br>F-ratio,<br>*p*-value |
| --- | --- | --- | --- | --- | --- | --- | --- | --- |
| SMR (mg/kg/h) | Quantile<br>(0.1) | 52.75 ±<br>1.27 | 54.76 ±<br>1.30 | 53.57 ±<br>1.31 | 47.82 ±<br>2.58 | - | - | - |
| | Quantile<br>(0.15) | 53.23 ±<br>1.22 | 55.50 ±<br>1.24 | 54.12 ±<br>1.48 | 48.42 ±<br>2.73 | - | - | - |
| | Quantile<br>(0.2) | 53.68 ±<br>1.03 | 56.26 ±<br>1.45 | 54.74 ±<br>1.60 | 48.71 ±<br>2.77 | 0.18, 0.68 | 5.06, **0.039** | 3.24, 0.066 |
| | Quantile<br>(0.25) | 53.87 ±<br>1.03 | 56.65 ±<br>1.40 | 55.25 ±<br>1.71 | 49.17 ±<br>2.87 | - | - | - |
| Net area (mg/kg)* | GAM | 18.00 ±<br>8.26 | 72.02 ±<br>40.57 | 504.63 ±<br>97.08 | 652.11 ±<br>97.11 | 8.11, **0.014** | 5.38, **0.037** | 3.29, 0.093 |
| SDAendtime (min) | GAM | 253.09 ±<br>97.42 | 737.33 ±<br>305.01 | 2323.45 ±<br>191.17 | 2668.64 ±<br>201.29 | 79.17, **<0.001** | 1.61, 0.22 | 0.10, 0.75 |
| MRpeak<br>(mg/kg/h)* | GAM | 62.55 ±<br>2.90 | 71.50 ±<br>1.49 | 80.11 ±<br>6.95 | 82.96 ±<br>7.64 | 12.69, **0.0024** | 0.0026, 0.96 | 0.036, 0.85 |
| MRfactscope | GAM | 1.17 ±<br>0.07 | 1.27 ±<br>0.03 | 1.45 ±<br>0.08 | 1.73 ±<br>0.20 | 10.96, **0.0044** | 1.36, 0.26 | 0.90, 0.36 |
| Net TAN excretion<br>(mmol/kg) † | - | 0.74 ±<br>0.15 | 0.98 ±<br>0.19 | 0.98 ±<br>0.13 | 1.36 ±<br>0.16 | 2.37, **0.030** | 3.10, **0.013** | 0.39, 0.54 |

\*trial *p*<**0.05**, † adjusted *p*-values by least squares regressions. GAM = General Additive Model.

**Figure 2.**
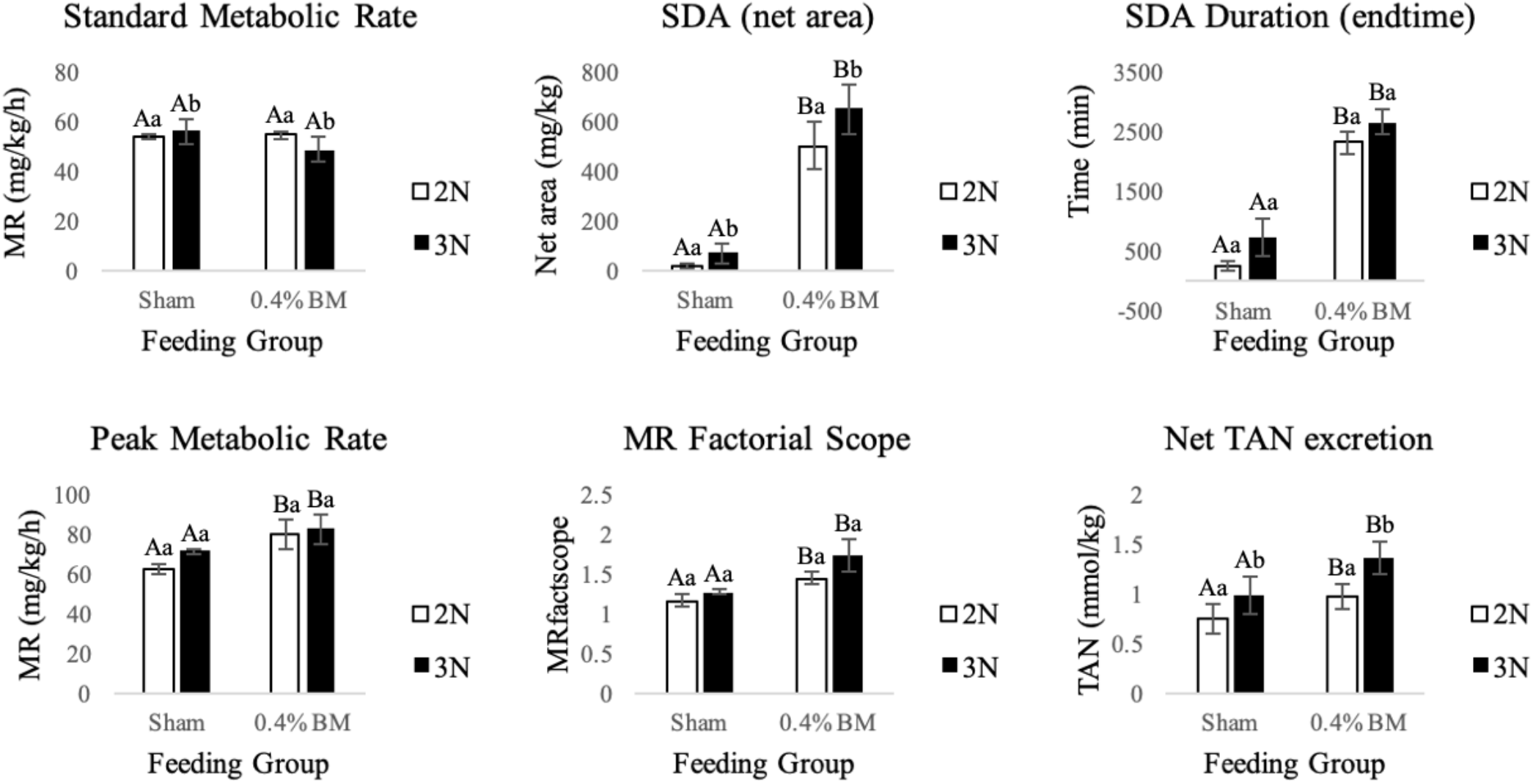
Graphed results of multiple-factor ANOVAs or ANCOVAs on the effect of feeding group and ploidy on standard metabolic rate (SMR), specific dynamic action (SDA) parameters and net total ammonia nitrogen (TAN) excretion in brook charr, *Salvelinus fontinalis* (mean ± SE). Parameters with different uppercase letters represent a significant feeding group effect, while lowercase letters represent a significant ploidy effect (*p*<0.05). For SMR and SDA parameters, n=4, 4, 6, and 6 for 2N-Sham, 3N-Sham, 2N-0.4% body mass (BM) and 3N-0.4% BM, respectively, and for net TAN excretion, n=7, 7, 9, and 9, respectively. Statistically significant *p*-values have been bolded (*p*<0.05).

Ploidy had a significant effect on SMR (triploids lower than diploids), SDA (net area; triploids higher than diploids), and net TAN excretion (triploids higher than diploids) (Table 2; Figure 2). The interaction between feeding group and ploidy was not significant for any of the measured variables and therefore no post-hoc tests could be completed to further investigate these trends.

## 4. Discussion

This study examined how triploidy affects standard metabolic rate and postprandial metabolism (including ammonia excretion) in fish, using brook charr as the model organism. This was done by feeding previously fasted diploids and triploids a single meal and then measuring their metabolic and excretory responses in comparison to their pre-feeding states. When combining data for sham-fed controls and fed fish, triploids had significantly lower SMRs, significantly higher postprandial MO_2_ (SDA net area), and significantly higher net TAN excretion values than their diploid counterparts. All parameters of SDA and net TAN excretion were significantly higher in the fed groups than in the sham-fed controls.

Body mass, fork length, and condition factor were not significantly affected by feeding group, ploidy, or their interaction. However, relative organ sizes (VSI, HSI, and GSI) were all significantly lower in triploids than in diploids. As oocyte development is blocked prior to vitellogenesis in triploids (Benfey 1999), significantly lower GSIs were anticipated for females; this likely explains the overall reduction in GSI, even though both sexes are represented in these data. Lower HSIs in triploids have also been documented in rainbow trout (Fauconneau *et al.* 1990) and Nile tilapia (*Oreochromis niloticus;* Hussain *et al.* 1995), but Segato *et al.* (2006) found no differences in HSI when comparing ploidies of shi drum. The decreased HSI in triploids likely stems from the lack of hepatic vitellogenin production in females (Benfey *et al.* 1989; Fauconneau *et al.* 1990; Hussain *et al.* 1995). Hepatic protein metabolism is highly responsive to feeding in diploid rainbow trout (McMillan *et al.* 1992), and the level of responsiveness could potentially be affected by ploidy. The lower VSI in triploids in the current study was due to the lower relative mass of the gonads and liver; this ploidy-related difference was no longer present when subtracting gonad and liver mass from the total visceral mass (data not shown). The categorical random variable trial was significant in the analyses of SDA and its peak MO_2_ (net area and MRpeak, respectively).

The effect of ploidy on SMR in this study (triploids lower than diploids) is opposite to that found by O’Donnell *et al.* (2017), although in the latter case the ploidy effect was not statistically significant. The study by O’Donnell *et al.* (2017) was completed at 16.2-17.4°C, which is higher than the thermal optimum for diploids (15°C; Smith and Ridgway 2019). Therefore, MO_2_ was likely elevated by thermal stress, and probably more so in the triploids based on results from other studies (Hyndman *et al.* 2003a; Atkins and Benfey 2008; Sambraus *et al.* 2017a; Sambraus *et al.* 2018). The lower temperature of the current study (13.7-15.0°C) may better reflect the effect of ploidy on SMR closer to the thermal optimum. The current study is the first to demonstrate significant differences in ammonia excretion between ploidies throughout SDA. Previous studies have compared ammonia excretion rates throughout early ontogeny in rainbow trout (Oliva-Teles and Kaushik 1987b) and post-exhaustive exercise in brook charr (Hyndman *et al.* 2003b), respectively, but without consideration of SDA.

As SMR is measured in a post-absorptive state and excludes periods of significant activity, differences in SMR likely stem from changes at the cellular level. Triploids are fundamentally different from diploids in this regard, having larger and fewer cells than diploids, and hence reduced cellular surface area to volume ratio and total cellular surface area for any given tissue (Benfey 1999). Such differences could affect basal metabolic processes through impacts on rates of respiratory gas transport (Schoenfelder and Fox 2015; Richardson and Swietach 2016) and the accumulation of older and damaged (senescent) cells due to a slower turnover rate (Hansen *et al.* 2015; Lahnsteiner and Kletzl 2018), perhaps due to reduced basal titres of chaperone proteins (Saranyan *et al.* 2017). While having fewer cells to maintain/turnover may explain the lower SMR of triploids in the current study, an increase in senescent cells could limit triploids metabolically when situations of high oxygen demand arise (e.g., Altimiras *et al.* 2002; Verhille *et al.* 2013). However, many studies have found no ploidy-related differences in maximum metabolic rate (Sezaki *et al.* 1991; Parsons 1993; Bernier *et al.* 2004; Lijalad and Powell 2009; Bowden *et al.* 2018).

Some previous studies suggest that triploids have a lower absolute or factorial aerobic scope compared to diploids (Altimiras *et al.* 2002; Bernier *et al.* 2004; O’Donnell *et al.* 2017). Consequently, they may be aerobically constrained when consuming large meal sizes relative to their BM, which could be mitigated by consuming smaller meals on a more frequent basis. However, other studies have found no effect of triploidy on aerobic scope (Sezaki *et al.* 1991; Parsons 1993; Lijalad and Powell 2009; Bowden *et al.* 2018). Future studies should track the aerobic scope of diploids and triploids throughout their lives to determine whether ploidies become more/less similar as they develop.

Lower protein reserves could explain the increased net TAN excretion that was observed in triploids, as ammonia is an end-product of protein metabolism. Segato *et al.* (2006) found that triploid shi drum had significantly lower retention (storage) of crude protein compared to diploids. Differences in HSI in the current study also support the possibility that triploidy affects protein metabolism. Due to the regular feeding intervals (of high protein diets) in aquaculture, these differences may not influence triploid performance. Nevertheless, diets with higher lipid or carbohydrate levels should be considered to potentially enhance energy storage and reduce the waste of dietary protein. Sacobie *et al.* (2016) found that lipid digestibility was unaffected by ploidy in brook charr; these findings, in tandem with the current study, suggest that triploids may be better suited for high lipid diets.

Although postprandial MO_2_ (SDA net area) was significantly higher in triploids, specific (pairwise) trends could not be investigated because the interaction between feeding group and ploidy was not significant. All other measured metabolic parameters were similar between ploidies, apart from the lower SMRs in triploids. However, triploids tended to have greater variability in their post-feeding MO_2_ (SDA) among individuals than the diploids (visualized in Figure 1). Triploid variability in metabolic rate was also documented in white crappies (*Pomoxis annularis*) by Parsons (1993). The combination of increased metabolic variability and a lower SMR may contribute to the higher SDA net area of triploids. If triploids invest more energy into postprandial metabolism, then they will likely be constrained metabolically while feeding under suboptimal conditions (e.g., high temperature, hypoxia, and exhaustive exercise). Further studies to investigate this hypothesis are warranted. Future research should also focus on TAN excretion; although the current study gave an accurate subsample of net TAN excretion, it was unable to capture the true total TAN excretion throughout SDA. Additionally, future studies should focus on protein synthesis rates to further investigate the physiological processes underlying differences in postprandial metabolism between ploidies.

## Acknowledgements

This study was completed in partial fulfillment of the requirements for a Master’s degree in Biology at UNB (NJD) and was supported financially by the Natural Sciences and Engineering Research Council of Canada (TJB Discovery Grant 2017-04397), the New Brunswick Innovation Fund (NBIF) (TJB Graduate Research Assistantship 2018-015), and the NBIF Science, Technology, Engineering, and Mathematics Award (NJD; 2017). We would like to thank members of the Benfey and Sacobie labs for their assistance with sampling.

## Notes

### Competing Interest Statement

The authors have declared no competing interest.

